# Hyperpolarization-Activated Currents Drive Neuronal Activation Sequences in Sleep

**DOI:** 10.1101/2023.09.12.557442

**Authors:** Dhruv Mehrotra, Daniel Levenstein, Adrian J Duszkiewicz, Sofia Skromne Carrasco, Sam A Booker, Angelika Kwiatkowska, Adrien Peyrache

## Abstract

Sequential neuronal patterns are believed to support information processing in the cortex, yet their origin is still a matter of debate. We report that neuronal activity in the mouse head-direction cortex (HDC, i.e., the post-subiculum) was sequentially activated along the dorso-ventral axis during sleep at the transition from hyperpolarized “DOWN” to activated “UP” states, while representing a stable direction. Computational modelling suggested that these dynamics could be attributed to a spatial gradient of hyperpolarization-activated current (I_h_), which we confirmed in *ex vivo* slice experiments and corroborated in other cortical structures. These findings open up the possibility that varying amounts of I_h_ across cortical neurons could result in sequential neuronal patterns, and that travelling activity upstream of the entorhinal-hippocampal circuit organises large-scale neuronal activity supporting learning and memory during sleep.

**Highlights:**

- **Neuronal Activation Sequence in HDC**: neuronal activity was sequentially reinstated along the dorsoventral axis of the HDC at UP state but not DOWN state onset.
- **Role of I_h_ in Sequence Generation**: Incorporating the hyperpolarization-activated current (I_h_) into computational models, we identified its pivotal role in UP/DOWN dynamics and neuronal activity sequences.
- ***Ex Vivo* Verification**: slice physiology revealed a dorsoventral gradient of Ih in the HDC.
- **Implications Beyond HDC**: the gradient of I_h_ could account for the sequential organization of neuronal activity across various cortical areas.

## Introduction

Neuronal computations arise from the coordination of neural activity across a wide range of spatiotemporal scales, from precisely timed action potentials in local circuits^1–6^ to coordinated fluctuations of neuronal activity over multiple brain areas^7–10^. This is particularly the case during spontaneous activity such as sleep, when computational functions must be carried out by internally organised activity patterns that constrain and coordinate neuronal spiking across thalamocortical networks^11–14^.

During the non-Rapid Eye Movement (NREM) stage of sleep, populations of cortical neurons alternate between hyperpolarized DOWN states and depolarized UP states with low-rate asynchronous spiking^15^. This “slow oscillation” originates in the neocortex^16^, entrains activity in thalamic and hippocampal circuits^8,17–19^, and is essential for learning and memory^20^. UP/DOWN alternations are thought to emerge from the interplay between connectivity and intrinsic neuronal properties^21,22^, and are well-fit by a model in which the UP state is maintained by recurrent excitation while the DOWN state is terminated by an adaptive process^23^. However, the biophysical identity of this adaptive process is unknown.

In local networks, individual neurons show sequential activation at the time of UP state onset^5,24^. This reliable sequence of spike timing provides a backbone against which perturbations represent specific stimuli^25^ and implement sleep-dependent functions^26,27^. Despite its functional significance, the mechanism that determines spike timing is unknown. It has been suggested that UP onset sequences reflect network-level properties of the cortex, such as connectivity^5^, or intrinsic differences between cells, such as excitability^24^. Thus, the mechanism of spike timing at the UP state onset is closely related to the generating mechanism of UP/DOWN states^22,28^.

To address the questions of the origin of sequential activity at UP state onset, we investigated population activity in the HDC during wakefulness and NREM sleep. The behavioural correlates of HDC neurons are well established, with a majority of principal neurons coding for the animal’s head direction during wakefulness^29^, and maintaining a coherent representation of a “virtual” head direction during sleep^14,30^. HDC extends dorso-ventrally, where the dorsal portion is proximal to the retrosplenial cortex and the ventral portion is proximal to the subiculum, and it is possible to record *in vivo* along the entire extent of its primary anatomical axis which allows the observation of a large number of neurons in the same area. Finally, the HDC is a critical hub of the entorhinal-hippocampal formation, where its activity during sleep plays a key role in memory processes.^20,31^. The HDC is thus an ideal model to investigate the organisation of neuronal activity on the population level.

Here, we report that spike sequences at the UP onset are spatially organized along the HDC dorsoventral axis, a phenomenon that was reproduced *in silico* with a gradient of I_h_ and verified in *ex vivo* patch clamp experiments. Finally, we show that variation in I_h_ can account for sequential activity at UP state onset in other cortical structures beyond HDC, suggesting that this intrinsic neuronal property plays a key role in generating and timing cortical dynamics.

## Results

We implanted linear silicon probes to target the entire dorsoventral extent of the HDC along a given layer (**Figure 1A**, **Supp Fig 1A**, Methods), and recorded a total of 1657 single units, including 1097 putative excitatory cells (“EX”), in 16 sessions from 16 mice **(Supp Fig 1B-C)**. HD tuning, as well as every other behavioural correlate that was tested, did not show a gradient along the dorsoventral axis (**Figure 1B, Supp Fig 1D**). During NREM sleep, we observed widespread DOWN states (**Figure 1C**), detected as periods of low population firing (see Methods, **Supp Fig 1E**).

**Figure 1:**
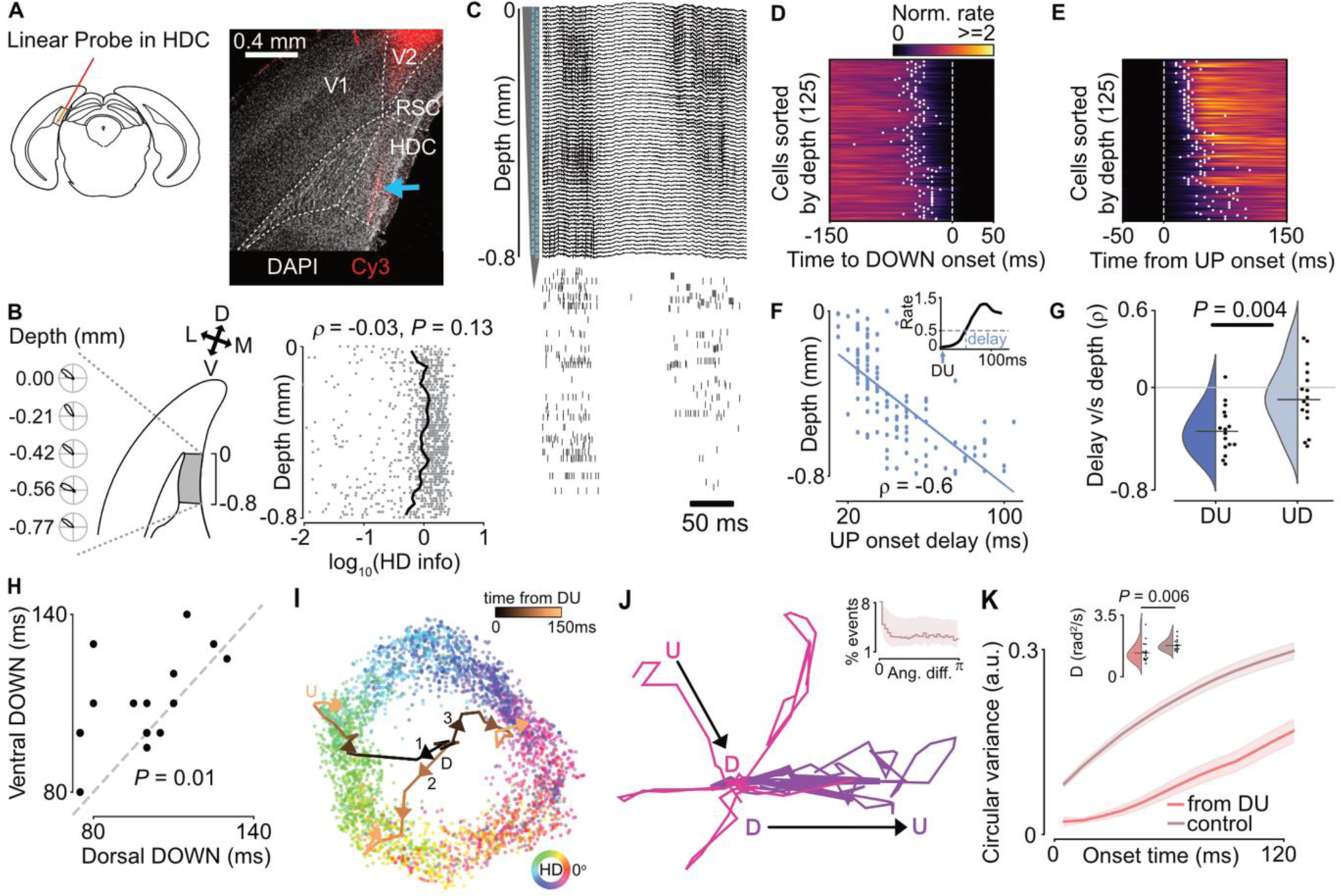
Locally generated sequential activation of HDC at the DOWN-to-UP transition. **A)** Left: Experimental schematic. Diagram adapted from the mouse brain atlas^32^. Right: Histology showing probe tract in HDC. **B)** Left: HD tuning curves as a function of HDC depth. Bottom: Quantification of HD information as a function of HDC depth. HD tuning is largely similar across the dorsoventral extent of HDC. (Kendall τ = -0.03, n = 1097, *p* = 0.13). **C)** Example UP and DOWN states observed in LFP as well as spiking. **D)** Peri-event time histogram (PETH) of cells aligned to the UP-to-DOWN (UD) state transition. White dots indicate the bin where the firing rate fell below 50% of the mean firing rate. **E)** Same as **D)**, but for DOWN-to-UP (DU) state transition. Note the sequential reinstatement of the UP state in the dorsoventral (top to bottom) direction as shown by the white dots indicating the bin where the firing rate exceeded 50% of the mean firing rate. **F)** Relation between UP onset delay and anatomical position for one example session (Kendall τ = -0.6, n = 125, *p* = 2.72 x 10^-21^). Inset: Threshold used to compute UP onset delay. **G)** UP onset delay v/s depth correlation for all sessions (Mann-Whitney U test, U = 52, n = 16, *p* = 0.004). **H)** DOWN state duration of dorsal versus ventral HDC for each session. Ventral DOWN states are longer than dorsal DOWN states (Wilcoxon signed-rank test, W = 9, n = 16, *p* = 0.01). **I)** Two-dimensional embeddings (using Isomap) of HD population activity during wakefulness (coloured points coded by angular direction) and trajectories on the manifold during three example travelling UP states (lines). Population activity tends to move in the same angular direction during the travelling UP state. **J)** Example decoded trajectories before and after the DOWN state, rotated by the angle at the first time bin where neuronal activity was sufficiently high for decoding (see Methods, **Supp Fig 1G**). Inset: Distribution of angular differences between the travelling UP state and the control. Bootstrapped confidence intervals (1000 shuffles of the angle at UD) given by the maximum and minimum values of the shuffled distribution for each bin are indicated by the shaded regions, suggesting that the observed distribution of angular difference is no different than a uniform distribution. **K)** Cumulative circular variance of the decoded angle during the travelling UP state (“from DU”) versus later in the UP state (“control”). Inset: Distribution of diffusion coefficients for all sessions, indicating that the angular change during the travelling UP state is overall lesser than later in the UP state, or more generally, during NREM (Mann-Whitney U test, U = 54, n = 16, *p* = 0.006).

### Neuronal activity at the DOWN-to-UP transition travels along the HDC dorsoventral axis and is locally generated

To examine the coordination of HDC population activity by the slow oscillation, we computed peri-event time histograms (PETHs) of the firing rate of all putative excitatory units aligned to the UP-to-DOWN (UD) and DOWN-to-UP (DU) transition. Unlike UD transitions, neurons activated with reliable delays with respect to the DU transition (**Figure 1D-F, Supp Fig 1F**), ranging from 5 ms to 100 ms (99^th^ percentile range), suggesting that they activated in a sequential manner. This sequential activity followed an anatomical pattern: dorsally located units tended to enter the UP state before ventrally located units (**Figure 1G**, One-sample Wilcoxon test for DU, W = 1, n = 16, *p* = 6.1 x 10^-5^ and for UD, W = 39, n = 16, *p* = 0.14). These neuronal activation sequences were not accounted for by differences in neuronal firing rates (**Supp Fig 1H**). Supporting this dorsalventral activation sequence, we found that the DOWN state durations were significantly longer in the ventral half, when detected separately from the dorsal half of HDC (**Figure 1H**, p = 0.013, Wilcoxon signed-rank test).

HD signal is coherent irrespective of brain state^14^. In order to characterize the dynamics of the HD representation in HDC during UP/DOWN transitions, we projected neural activity on a two-dimensional space using Isomap^33^, revealing a ring-shaped manifold that corresponded to real HD during wakefulness and ‘virtual’ HD during NREM ^30,34^ (**Figure 1I**). In this space, DOWN states were located at the centre of the ring, and DU transitions corresponded to a trajectory from the centre to the periphery of the ring^30,34^. We found that the DOWN state resets this virtual HD signal, as there was no relationship between the angle represented at the end of an UP state and the beginning of the subsequent UP state (**Figure 1J**). Following the DOWN state, the represented HD maintained a specific direction throughout the duration of the UP onset sequence (**Figure 1I**) and changed less rapidly than later in the UP state (**Figure 1K**).

We next sought to confirm the dorsoventral activity gradient in HDC with a measure independent of spike detection. DOWN states are accompanied by a signature pattern in the extracellular field potential (LFP) – a 0.5-4 Hz (delta) wave^35^ (**Figure 2A, Supp Fig 2A**) and drop in high gamma power (*γ*_h_: 70-150 Hz), a proxy for multi-unit activity^36^ not confounded by the firing rate, number, or quality of isolated units. We found that the delta waves (**Supp Fig 2B, C**) and drop in *γ*_h_ (**Figure 2B, C**) were synchronous across the DV axis at the UD transition. However, *γ*_h_ travelled in the DV direction during the UP state onset, mirroring the wave observed in single unit activity (**Figure 2B-F**, One-sample Wilcoxon test for DU, W = 21, n = 16, *p* = 0.013 and for UD, W = 68, n = 16, *p* = 1). The correlation coefficients between UP onset delay and depth for individual sessions were well-correlated between LFP and spikes (**Supp Fig 2D**) and the speed of the travelling activity for significant sessions was similar when computed using either the LFP or spiking activity (Spikes: 6.72 +/- 4.08 mm/s, LFP: 9.89 +/- 5.51 mm/s Mann-Whitney U test, U = 58, n = (13, 15), *p* = 0.072, **Supp Fig 2E**). Taken together, these findings reveal a dorsoventral sequence of neuronal activation at UP state onset in HDC, which is independently observed in both spikes and LFP.

**Figure 2:**
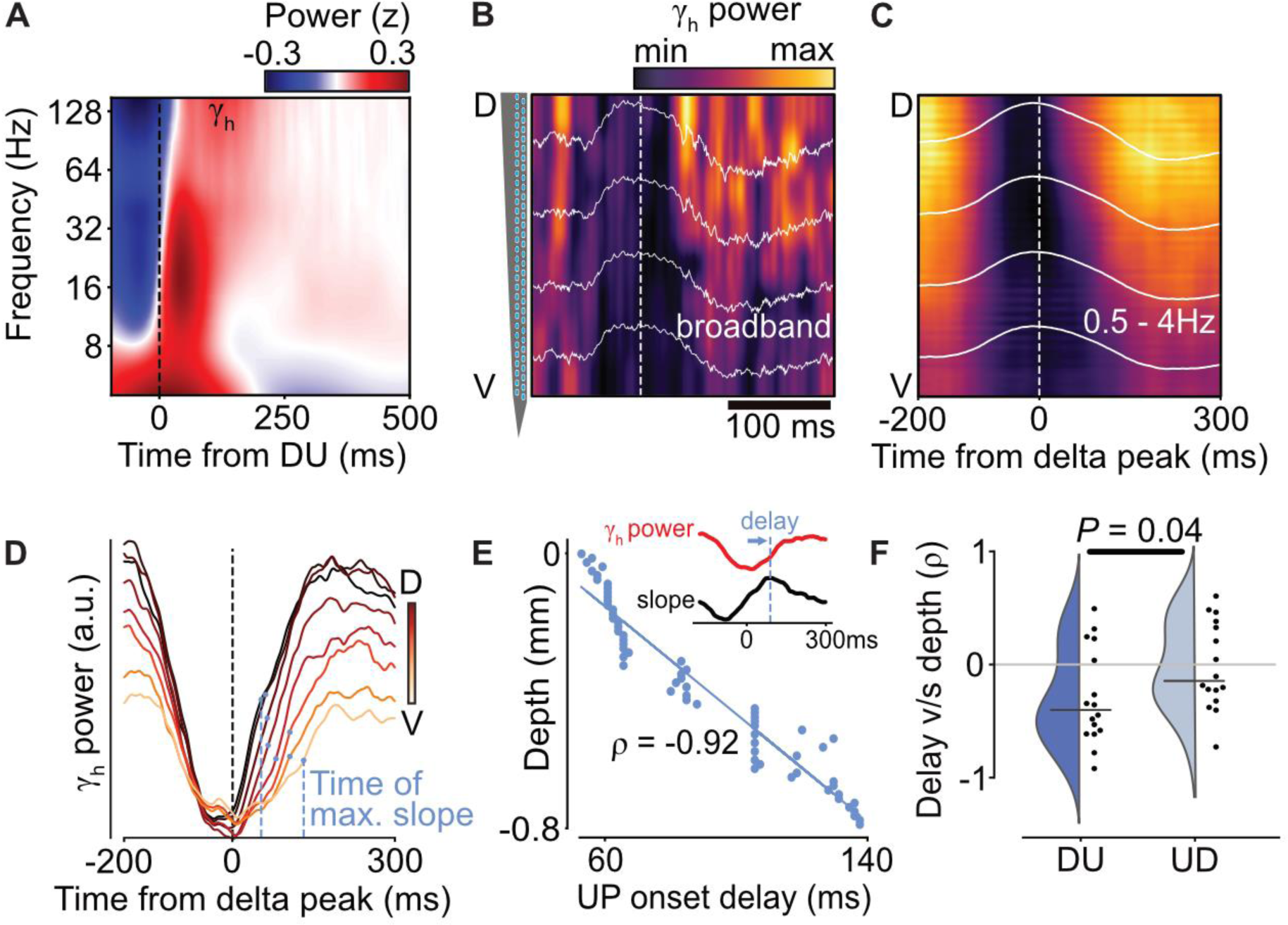
Gamma wavefront in HDC exhibits travelling activity. **A)** DU-triggered LFP spectrogram (average of epochs at least 500 ms in duration). Note increased power in the high gamma (*γ*_h_) band (70-150 Hz) right after DU. **B)** Example epoch showing broadband LFP (white, dashed line indicates the delta peak), and high gamma power post-delta. **C)** Same as **B)** but averaged for one session. Note the asymmetry in *γ*_h_ power reinstatement along the DV axis. **D)** Delta peak-aligned *γ*_h_ power for 8 example channels (coloured by anatomical position). Blue dots indicate the time of maximal slope for each trace. **E)** Relation between time of maximal slope and anatomical position for one example session (Kendall τ = -0.92, n = 64, *p* = 8.036 x 10^-26^). Inset: Illustration of time of maximal slope, referred to as “UP onset delay.” **F)** UP onset delay v/s depth correlation for all sessions (Mann-Whitney U test, U = 73, n = 16, *p* = 0.04).

The dynamics of neuronal activity at UP state onset can be inherited from their upstream inputs in the thalamus, where activity is also modulated by UP and DOWN states^37^. The HDC receives its primary inputs from the anterodorsal nucleus (ADN) of the thalamus, and activity in the two structures is strongly coupled across brain states, including NREM sleep^14,38^. To examine if there was any sequential activation during NREM sleep in the thalamus, we analysed ADN population activity during NREM from a previously published dataset^14^. At UP state onset, ADN cells activated more synchronously than in the HDC (**Supp Fig 2F, G**, standard deviation of delays: 10.38 +/- 5.74 ms in ADN vs 15.54 +/- 5.42 ms in HDC, Mann-Whitney U test, U = 29, n = (16, 9), *p* = 0.016). In other words, the highest variability in delays was observed at UP state onset in the HDC, suggesting that the dorsoventral sequential activation is the consequence of a local property of the HDC.

### An I_h_ gradient can account for dorsoventral sequential activation at UP state onset

To elucidate the mechanisms behind the sequential activation at the UP state onset, we modelled UP/DOWN alternation dynamics using an Adapting Wilson-Cowan (AdWC) model^23^ (**Figure 3A**), which captures the alternation between UP and DOWN states via an interplay between recurrence, fluctuating input, and an adaptive process.

**Figure 3:**
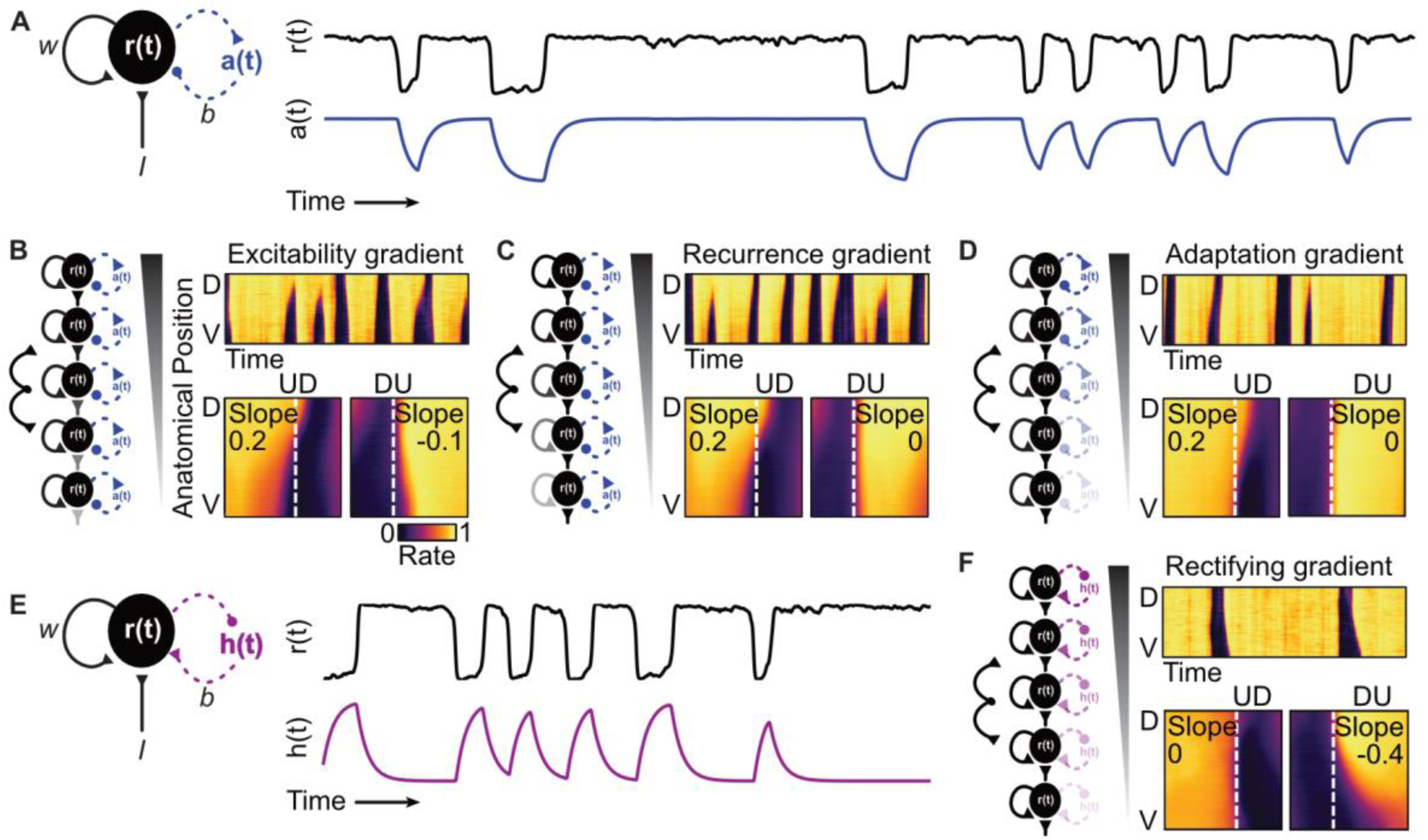
An I_h_ gradient can account for the travelling wavefront at UP state onset only. **A)** Left: Schematic of the adapting Wilson-Cowan model. Right: Example simulation of UP and DOWN alterations generated by this model. Adaptation *a(t)* is high when the population rate *r(t)* is high. **B-D)** Biophysical parameter gradients in excitability **(B)**, recurrence **(C)**, and adaptation **(D)** in a linear array of the adapting Wilson-Cowan model are unable to replicate experimental observations. In each panel, top right: Example simulation of population rates. Bottom right: Average PETH of the simulation. **E)** Left: Adapting Wilson-Cowan model now with a rectifying current *h(t)* (thought to be functionally equivalent to I_h_) instead of adaptation. Right: Example simulation of UP and DOWN alterations generated by this model. Rectifying current is high when the population rate is low. **F)** A gradient of rectifying currents can replicate the experimental observations; observe how the UD transition is synchronous while the DU transition travels dorsoventrally uniquely in this simulation.

We hypothesised that the sequential activation in HDC could arise from a gradient of a biophysical parameter along the DV axis and modelled a linear array of AdWC units with a gradient in either excitability, the degree of recurrent excitation, or the strength of an adaptive current (Methods, **Figure 3B-D**). Surprisingly, we found that none of these parameter gradients were able to account for our primary observation: synchronous deactivation at the UD transition, but sequential activation at UP state onset (**Figure 3B-D**). All simulations were tested with a wide range of parameters (**Supp Fig 3B-D**), yet no gradient in these parameters was able to replicate the experimental observations.

The intuition behind these results is as follows: because each of the parameters affect the stability of the UP state, and the UD transition is attributable to perturbations out of the UP state, these gradients in turn have a strong effect on the timing of the UD transition (**Figure 3B-D**). This suggests that the biophysical property underlying the experimental observation must come into effect exclusively during the DOWN state, thus only influencing the timing of the DU transition. One such biophysical property is the hyperpolarization-activated current (I_h_): a depolarizing current activated by hyperpolarized voltages and prevalent in cortical pyramidal cells^39,40^.

Interestingly, I_h_ is functionally similar to adaptation in the Wilson-Cowan formalism, as it can be implemented as an additive current with an inverse activation profile to adaptation (**Supp Fig 3E**). Indeed, adding a uniform I_h_ to the model was able to produce UP/DOWN alterations (**Figure 3E**). However, when a gradient of I_h_ was included, it had a distinct signature from adaptation, due to its differential effect during the DOWN, rather than the UP state. As a result, an I_h_ gradient led to synchronous UD and sequential activation at UP state onset, as observed in our recordings (**Figure 3F**). Importantly, this property of the network was robust to a wide range of parameters (**Supp Fig 3F**).

We next sought to verify the presence of an I_h_ gradient in the HDC. We performed whole-cell patch clamp recordings in layer III excitatory cells along the DV axis of the HDC (**Figure 4A, B**), and tested for the presence of a gradient of hyperpolarization-activated cyclic nucleotide gated (HCN)-channels, which are believed to mediate I_h_ currents^41–44^. I_h_ was quantified as the relative magnitude of “sag” potential after hyperpolarizing the neuron^45–48^ (**Figure 4C**). We found a gradient of decreasing “sag” along the DV axis (**Figure 4D**) which was sensitive to an HCN-channel blocker (20 μM ZD-7288) (**Figure 4C, D**). Critically, no other dorsoventral gradients in passive properties or excitability were found in HDC cells **(Supp Fig 4)**. Together, these results suggest that sequential activation at UP state onset can be attributed to a gradient of I_h_.

**Figure 4:**
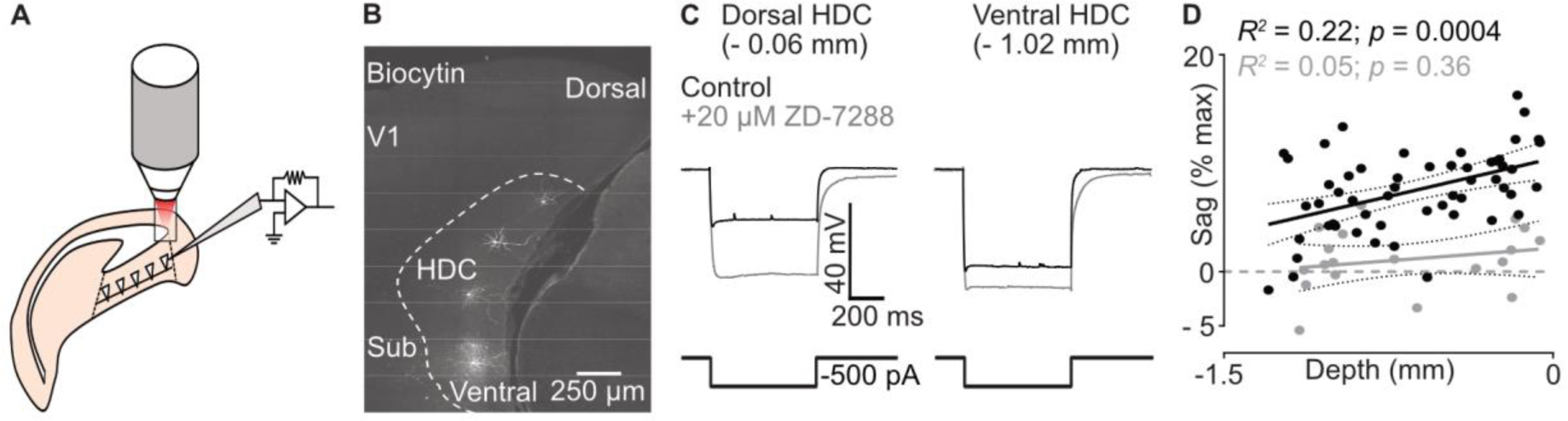
Theoretically predicted I_h_ gradient observed *ex vivo* in HDC. **A)** Experimental schematic. **B)** Representative microscopy overview of the recorded cells labelled with Biocytin. V1 = Primary visual cortex, Sub = Subiculum. **C)** Representative voltage responses of dorsal and ventral HDC cells to a hyperpolarizing pulse (black), and in the presence of HCN channel blocker ZD-7288 (grey). The voltage “sag” is diminished in the presence of ZD-7288. **D)** Quantification of voltage “sag” response as a function of the anatomical position of recorded cells; a dorsoventral gradient in “sag” is observed along the HDC (black, Pearson R^2^ = 0.22, n = 59 cells from 11 mice, *p* = 0.004), which is abolished in the presence of ZD-7288 (grey, Pearson R^2^ = 0.05, n = 19 cells from 11 mice, *p* = 0.36).

### Post-DOWN rebound suggests a widespread role for I_h_ in neuronal activation at UP onset

In addition to earlier activation at UP state onset, our model predicts that units with strong I_h_ should have a greater firing rate rebound compared to their late-activating, low-I_h_ counterparts (**Figure 5A**).

**Figure 5:**
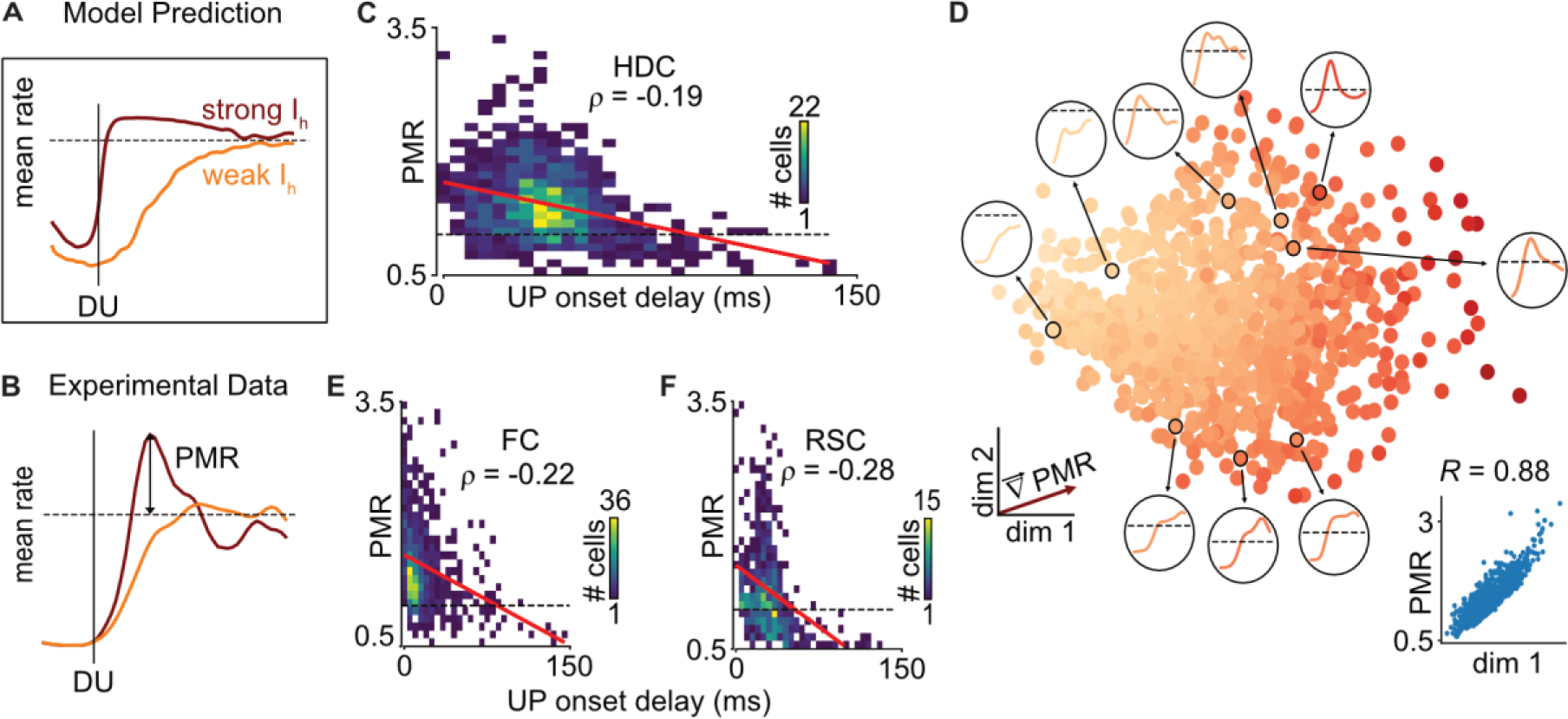
Post-DOWN rebound suggests a widespread role for I_h_ in UP onset timing. **A)** A prediction of the I_h_ gradient is that cells with stronger I_h_ would show an increase in firing rate overshoot (i.e., PMR) from the mean at DU (brown), compared to cells with weaker I_h_ (orange). **B)** Example cells from experimental data with low and high PMR (orange and brown, respectively). **C)** High PMR units exhibit earlier UP onset, suggesting that PMR as a measure of I_h_ can be used to predict when a given unit enters the UP state. This general principle was found to hold in HDC (Kendall τ = -0.19, n = 1096, *p* = 3.98 x 10^-20^). **D)** Two-dimensional Isomap projection of PETHs aligned to DU (+/- 250ms). Points represent PETHs of individual HDC cells, coloured by PMR (darker colours indicate larger PMR). The projection is surrounded by representative PETHs (in black circles), the dashed line indicating the mean firing rate of the cell. Left inset: Gradient vector of the Isomap projection space indicating the direction of most variance in PMR. The magnitude of the gradient vector indicates that the first dimension of Isomap accounts for most of the variance in the data. Right inset: First dimension of Isomap projection correlates well with PMR (Pearson R = 0.88, n = 1096 EX units from 16 sessions (16 mice), *p* = 0). **E-F)** Same as **D)**, but for frontal cortex (FC, data from^49^, Kendall τ = -0.22, n = 950 EX units from 25 sessions (11 rats), *p* = 5 x 10^-21^) and **F)** retrosplenial cortex (RSC, Kendall τ = -0.28, n = 563 units from 14 sessions (3 mice), *p* = 6.76 x 10^-21^)

We therefore measured the peak firing rate of each unit at UP state onset, normalised to its mean rate during the UP state (i.e., “peak-to-mean ratio,” PMR, **Figure 5B**). As predicted, PMR was negatively correlated with UP state onset timing, with earlier-spiking cells showing a larger overshoot (**Figure 5C**). Critically, virtually no units with a late activation at UP onset had a large PMR. Further, we found that PMR could account for a large part of the variance of spiking activity at the UP state onset. Isomap decomposition of the DU-PETHs of all HDC units (**Figure 5D**) revealed a continuous gradation of temporal profiles at the UP state onset, which was largely attributable to a single dimension strongly correlated with PMR (Pearson R = 0.88, n = 1096, *p* = 0, **Figure 5D**, inset, **Supp Fig 5**).

We then examined if the relationship between PMR and UP onset delay was found in cortical regions other than HDC. We found a similar relationship in the frontal cortex (FC, data from^49^) (**Figure 5E**) and retrosplenial cortex (RSC, **Figure 5F**). Overall, these observations suggest that cortical sequences at the UP state onset have an intrinsic origin (not only in the HDC, but also in other regions) namely, variability in the strength of hyperpolarization-activated currents (I_h_).

## Discussion

By recording the HDC with linear silicon probes, we have shown that neurons are sequentially activated from dorsal to ventral HDC at the DOWN to UP state transition. This sequential activation was locally generated at UP state onset, as the transition to DOWN state was synchronous and neurons in the HDC’s primary thalamic input showed a narrower distribution of reinstatement delay. A mathematical model of cortical activity revealed that these experimental observations could be accounted for by a gradient in the strength of a hyperpolarization-activated rectifying current (I_h_). This gradient of I_h_ was confirmed by *ex vivo* patch clamp experiments, and successfully predicted that earlier reinstatement at UP state onset was associated with an overshoot of activity. We confirmed this relationship in the HDC and other cortical areas. These results suggest that sequential activity during NREM sleep has an intrinsic origin in hyperpolarization-activated currents and have implications for memory formation in the downstream entorhinal-hippocampal network.

### Activity patterns during NREM sleep have an intrinsic origin in hyperpolarization-activated currents

The slow oscillation is the most prominent neocortical activity pattern during NREM sleep^15,35^, which consists of aperiodic transitions from a depolarized UP state to transient hyperpolarized DOWN states. The exact mechanism behind the slow oscillation is not well understood, but several candidate mechanisms have been proposed including Na^+^-dependent K^+^ current activation^22^, synaptic fatigue^28,50^, and GABA_B_ activation^51^. In each of these cases, the slow oscillation is hypothesised to rely on the interplay between recurrent excitation, which maintains activity during the UP state, and a slow adaptive process, which is activated by neuronal activity during the UP state and disengaged by neuronal inactivity during the DOWN state. Models using these principles can produce realistic UP/DOWN alternation dynamics, including *in vivo*-like duration statistics^23,52^ and spatially travelling waves^22,53^. Indeed, the slow oscillation has been observed to travel across cortical layers^16,54^ and laterally within and across cortical areas^5,9,55–57^. However, the travelling UP onset we observed is different from these previous reports in that the DOWN onset (and delta wave) itself does not travel, but only the return to UP state does. We took advantage of this unique phenomenon to unravel a novel mechanism of slow wave generation in the HDC.

Our computational modelling suggests that the adaptive process responsible for timing the cortical slow oscillation in HDC is the hyperpolarization-activated current (I_h_). Where traditional adaptive processes provide slow negative feedback on neuronal activity, the net effect of I_h_ is positive feedback on neuronal inactivity, which can play an analogous role in the generation of the slow oscillation but has distinct implications for its dynamics and accounts for biological observations. First, adaptation is only expected to affect high-rate neurons, yet cortical neurons spike at low rates *in vivo*^49^. As all neurons are hyperpolarized during DOWN states, I_h_ affects all neurons irrespective of their activity rate. Second, the variable duration of low-rate UP states *in vivo* suggests a stable UP state with balanced excitation and inhibition^52,58^. In order to produce *in vivo*-like duration statistics, an adaptation-based mechanism would require tonically active adaptation throughout the duration of the UP state^23^. In our model, the activation of I_h_ ends an otherwise-stable DOWN state and intrinsic differences in the strength of I_h_ determine the relative timing of neuronal reinstatement after the DOWN state.

Our model using an inactivity-based adaptive process made another prediction, irrespective of the spatially structured activation of neurons specific to HDC: neurons quickly reinstating their activity should show an overshoot at UP state onset, while slowly reinstating neurons should monotonically increase their activity to their basal firing rate. We have confirmed this prediction across different datasets from three cortical structures. Hence, I_h_ may be a key factor accounting for rich dynamics of cortical activation at UP state onset^5^. Importantly, neurons should consistently show the same temporal profile at UP state onset, only depending on intrinsic currents and independent of the circuit connectivity^5^ or intrinsic excitability of the neurons^24^.

I_h_ is mediated by hyperpolarization-activated cyclic nucleotide-gated (HCN) ion channels, which are known to be important in the generation of spontaneous and oscillatory activity in several brain regions^59–64^. Here, we reveal that I_h_ may be a ubiquitous mechanism for slow wave generation in cortical tissue and it may play a role in population-level dynamics at UP state onset.

### Implications for memory formation during sleep in the entorhinal-hippocampal network

The entorhinal-hippocampal network is organised along a principal axis corresponding to the dorsoventral axis of the entorhinal cortex and the septotemporal axis of the hippocampus^65,66^. This axis is characterised by gradients of intrinsic neuronal features^67,68^, size and spacing of spatial tuning^65,69–73^ as well as oscillatory properties at theta frequency^74–77^. It is believed that tuning and theta gradients result, at least in part, from a spatial gradient in I_h_^78,79^. Our observed lack of a spatial gradient of tuning properties in HDC poses the question of the significance of the I_h_ gradient and corresponding DV sequence during sleep for the navigation system.

The principal axis of the HDC maps onto the principal axis of the entorhinal-hippocampal formation^80^. While it has been suggested that the HDC codes for additional spatial features than an animal’s head-direction, for example speed^81^ and environmental boundaries^82^, these additional coding properties remain marginal, and the size of the HDC (especially relative to the ADN^83^) suggests a highly redundant code. One possibility is thus that the HDC is itself made of multiple modules distributed dorsoventrally which each provide head direction information for parallel loops within the entorhinal-hippocampal network. While the spatial gradient of I_h_ in HDC may not be sufficiently strong to affect tuning properties (**Supp Fig 1D**), I_h_ provides cells with specific computational properties^84–86^, for example determining their temporal integration window^87–90^ or resonance for input at specific frequencies^74–77^. Thus, another possibility is that the I_h_ gradient in HDC plays a role in integrating signals with different temporal profiles, such as angular head velocity signals conveyed from various parts of the visual system, or inputs differently modulated by theta (∼8 Hz) oscillations.

In the hippocampus, neuronal activity travels along its principal axis, both in wakefulness^91^ and in NREM sleep during sharp wave-ripples (SWRs)^92^. Our data suggest that sequential activity is also present upstream to the hippocampus and thus may play a role in the spatiotemporal organisation of SWRs. Although no direct inputs from HDC to the hippocampus has been described so far, this influence could be mediated through a disynaptic relay via the entorhinal cortex^93^.

During NREM sleep, hippocampal activity is locked to cortical slow waves^8,17,18,94^, which is believed to be instrumental for long-term memory formation^95,96^. The ADN is a crucial relay of cortical activity in orchestrating large-scale coordinated activity in the hippocampal and parahippocampal formation^34,97^. This thalamic nucleus mainly targets the HDC which, in turn, projects to the entorhinal cortex and other parahippocampal structures.

The role of the HD system for memory processes during sleep is still unclear, but one hypothesis postulates that it influences the downstream entorhinal-hippocampal formation by querying this network with specific directions^98^. Importantly, the sequential reinstatement of neuronal activity at UP state onset is faster than the typical drifting rate of the HD cell population during NREM sleep (**Figure 1I**). This suggests that the entire main axis of the entorhinal-hippocampal formation is sequentially activated at UP state onset with a barrage of excitation from HDC conveying a coherent and aligned head direction signal. Furthermore, the lack of relationship between the encoded angle at UP state onset and at the end of the previous UP state suggests that the DOWN state resets the activity in the HD network. Hence, each UP state would start from a new and possibly random direction, which would then continuously drift as long as the UP state is maintained. This has important implications for our understanding of the relationship between global brain dynamics (i.e., the slow oscillation) and information encoding in NREM sleep.

### Conclusion

Sequential activation of neurons is believed to play a key in cortical computation during rest^1,3,5,99^ and in many contexts beyond sleep^6,100,101^ . These sequences are generally thought to involve synaptic transmission^100^ or network effects^102^. However, our work introduces a novel potential mechanism for sequence generation in neural tissue – variable levels of I_h_ or of hyperpolarization could result in sequences that would not require precise synaptic transmission.

## Methods

### Animals

All procedures were approved by the Animal Care Committee of the Montreal Neurological Institute at McGill University in accordance with Canadian Council on Animal Care guidelines. The subjects were adult (> 8-week-old) mice bred by crossing wild-type females on C57BL/6J background (Jackson laboratories 000664) with homozygous male VGAT-IRES-Cre mice (Jackson laboratories 028862; n = 16). Mice were kept on a 12-hour light/dark cycle and were housed in group cages (2 – 5 mice per cage) before electrode implantation surgery and individually afterwards.

For the RSC dataset, the subjects used were adult C57BL6/J mice (n = 1) and mice bred by crossing wild-type females on C57BL/6J background (Jackson laboratories 000664) with homozygous male FVB/NJ mice (Jackson laboratories 001800; n = 2).

*Ex vivo* electrophysiological experiments were performed in accordance with institutional (University of Edinburgh, UK) and UK Home Office guidelines. Mice were maintained on a 14-hour light/dark cycle, housed in group cages (4 – 6 mice per cage), and given *ad libitum* access to food and water. Experiments were performed in acute brain slices prepared from 8– 12-week-old male C57/Bl6JCRL mice, which were either bred in house or purchased directly from Charles River, UK.

### Electrode implantation

C57BL/6 mice (Jackson Laboratory) were implanted under isoflurane anaesthesia, as previously described in^103^. Silicon probes were mounted on in-house build movable microdrives and implanted through a small craniotomy. Probes were implanted at a 26-degree angle pointing away from the midline towards the HDC (from Bregma: AP, -4.24 mm; ML, 1.70 mm; DV, -1.00 mm). A mesh wire cap was then built around the implanted microdrive and was reinforced with UV-cured adhesive. Mice were allowed to recover for at least 3 days before electrophysiological recordings. The probes consisted of a single shank with 64 recording sites (H5, Cambridge Neurotech, electrodes staggered in two rows, 25 μm vertical separation). In all experiments both ground and reference wires were soldered to a single 100 μm silver wire which was then implanted 0.5 mm into the cerebellum.

For RSC recordings, procedures were similar in all aspects. Probes were implanted into the left RSC (from Bregma: AP, -3.1 mm; ML, -1 mm; DV, -0.3 mm). One of the RSC animals was implanted with a probe having 8 shanks with 64 recording sites (Buzsaki64, Neuronexus, electrodes staggered in 2 rows, 8 recording sites per shank, 200 μm shank spacing, 20 μm recording site spacing), and tungsten wires in left hippocampal CA1. This animal was implanted at an angle from midline, more or less parallel to the RSC layers (from Bregma: AP, -3.7 to -2.29 mm; ML, -1.1 to -0.5 mm; DV, -0.5 mm).

### Recording procedures

During the recording sessions, neurophysiological signals were acquired continuously at 20 kHz on a 256 channel RHD USB interface board (Intan Technologies). The wide-band signal was downsampled to 1.25 kHz and used as the LFP signal. The animals were tethered to a motorised electrical rotary joint (AERJ, Doric Lenses) to enable free movement around the enclosure. Ahead of the main recording session, the microdrive was lowered over several hours in small (35 - 70 μm) increments until the whole shank was positioned in HDC. The dorsal edge of HDC was inferred from the sudden appearance of single units narrowly tuned to HD, and the shank was positioned so that these HD-tuned units were present on the top-most channels^103^. For the animals included in the analysis (n = 16), the probe tip was advanced an average of 2.04 mm into the brain tissue (SD = 0.19 mm), with the top channel of the probe at the depth of 1.24 mm. One animal was excluded from the analysis since the probe tip was advanced 2.54 mm into the brain tissue before HD-tuned units were observed at the bottom channels, making it unlikely that the recording spanned the dorsal half of the HDC.

A short open field session was then recorded to map the HD receptive fields of all neurons. The recording depth was adjusted so that sharply tuned HD cells (a hallmark of the HDC) are present along the entire length of the shank. Data collection did not commence until at least 2 hours after the last depth adjustment. Animal position and orientation was tracked in 3D using 7 infrared cameras (Flex 13, Optitrack) placed above the enclosure and coupled to the Optitrack 2.0 motion capture system. Five small tracking markers were attached to the headcap and additional 2 larger markers were attached to the preamplifier chip. In addition, video recording was captured by an overhead camera (Flex 13, Optitrack) placed close to the rotary joint. Animal position and head orientation was sampled at 100 Hz (120 Hz for the RSC dataset) and was synchronised with the electrophysiological recording via TTL pulses registered by the RHD USB interface board (Intan). The tracking system was calibrated in the same manner across all recording sessions to ensure that the tracking coordinates were the same across the whole dataset.

### Behavioural procedures

Before the implant surgery, mice were habituated over several days to forage for small pieces of Honey Nut Cheerios cereal (Nestle) in the open field. For most recordings, the recording chamber consisted of a metal frame (90 x 90 x 180 cm) supporting a plastic platform with removable walls (width: 80 cm, height: 50 cm) that could be arranged into either a square or triangular open field. The recording protocol consisted of a sleep session in the home cage, followed by open field exploration in a square arena and another sleep session. A subset of animals then explored a triangular arena. A white rectangular cue card on one of the walls served as a salient cue. Both environments were oriented so that the wall with the cue card always faced the same direction.

For the RSC dataset, animals explored a circular arena (diameter 70 cm) with a white cue card, in addition to the square arena described above. A total of 563 units were recorded from 14 sessions.

## Tissue processing and probe recovery

Following the termination of the experiments, animals were deeply anaesthetised and perfused transcardially first with 0.9% saline solution followed by 4% paraformaldehyde solution. The microdrive was then advanced to remove the probe from the brain and the probe was moisturised with distilled water while the brain was being extracted. Brains were sectioned with a vibratome coronally in 40 μm thick slices. Sections were washed, counterstained with DAPI and Red Neurotrace and mounted on glass slides with ProlongGold fluorescence antifade medium. Sections containing probe tracts were additionally stained with a Cy3 anti-Mouse secondary antibody to help visualise the electrode tract. A widefield fluorescence microscope (Leica) was used to obtain images of sections and verify the tracks of silicon probe shanks. Probes were lifted out of the brain immediately after perfusion by turning the Microdrive screw all the way up. As the headcap was being manually dismantled, the probe shank was then kept from drying by infusions of distilled water into the headcap. Once the drive-mounted probe was separated from the headcap, it was immersed in 3% peroxide for 5 minutes and rinsed with distilled water. The probe was then dipped in and out of a warm Contrad solution for several minutes, followed by a 2-hour incubation in a warm 2% Tergazyme solution. The drive-mounted probe was then rinsed with distilled water and stored for the next implantation. This multistep cleaning protocol enabled us to implant an individual probe an average of 3 times.

### Spike sorting and unit classification

One recording session per mouse was included in the analysis. Spike sorting was performed semi-automatically, using Kilosort 2.0^104^ followed by manual curation of the waveform clusters using the software Klusters^105^. At this stage, any cluster without a clear waveform and clear refractory period in the spike train autocorrelogram were classified as noise and cluster pairs with similar waveforms and common refractory period in their spike train cross-correlogram were merged. Viable units were first identified as units that: (1) had an average firing rate of at least 0.5 Hz during open field exploration, and (2) had a waveform with negative deflection (criterion aiming to exclude spikes from fibres of passage). Next, putative excitatory cells and putative FS interneurons were classified based on their mean firing rate during the open field exploration and the through-to-peak duration of their average waveforms^103^. Putative FS interneurons were defined as cells with short trough to peak duration (<= 0.35 ms) and high mean firing rates (> 10 Hz). Conversely, cells with long trough-to-peak (> 0.35 ms) and low mean firing rates (< 10 Hz) were classified as putative excitatory (EX) cells. For the purposes of this study, only EX units were used for analyses.

### HD tuning curves and tuning metrics

The animal’s HD was calculated as the horizontal orientation of a polygon constructed in Optitrack from the 3-dimensional coordinates of all tracking markers, relative to the global coordinates which were constant across the whole study. HD tuning curves were computed as the ratio between histograms of spike count and total time spent in each direction in bins of 1 degree and smoothed with a Gaussian kernel of 3° s.d.

HD information contained in the tuning curves was calculated for N angular bins as:

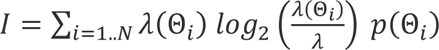

where λ(θ_i_) is the firing rate of the cell for the *i*th angular bin, λ is the average firing rate of the neuron during exploration and p(θ_i_) is the occupancy (i.e., normalised time spent) in direction θ_i_.

Linear speed scores and AHV scores were calculated by first binning spikes of each cell in 0.01 second bins and smoothing with a Gaussian kernel of 50 ms s.d. Linear speed and AHV were computed as horizontal/angular displacement of the animal in each tracking frame, divided by the sampling rate of 100 Hz. Speed score was defined as a Pearson correlation coefficient between the binned spike trains and the animal’s speed. AHV score was defined as the Pearson correlation coefficient between the binned spike trains and the animal’s AHV, and computed separately for clockwise and counterclockwise head turns. Both clockwise and counterclockwise AHV values were highly correlated with each other for most cells (data not shown), and the higher absolute value of the two was used as each cell’s AHV score.

### Classification of sleep states

Sleep scoring was performed using the SleepScoreMaster algorithm^49,106^ (available at https://github.com/buzsakilab/buzcode/tree/master/detectors/detectStates/SleepScoreMaster). The wide-band signal was down sampled to 1.25 kHz and used as the LFP signal.

### UP and DOWN state detection

UP and DOWN states were detected as previously described^34^. The total firing rate of all simultaneously recorded neurons in bins of 10 ms was computed and smoothed with a Gaussian kernel of 20 ms s.d. Epochs during which total firing rate was lower than the 20^th^ percentile of the population rate was considered DOWN states. Epochs shorter than 30 ms and longer than 500 ms were discarded. UP states were defined as the epochs between each DOWN state.

To compare the duration of DOWN states between dorsal and ventral HDC independent of the number and rate of the neurons in the two groups, (Figure 1H), we first normalized neuronal activity to the mean population rate and used a threshold of 0.2 (instead of the 20^th^ percentile of the population rate)

## Isomap Projection and Decoding

For Isomap analysis, only cells with HD information exceeding 0.2 bins/spike (see ref.^103^ for details) were included. Spike trains from all HD cells were binned during wakefulness (bin size of 200 ms), during periods when the animal had a speed of > 2 cm/s, and during NREM sleep (bin size of 50ms with 20% overlap). Different bin sizes were used to capture the different timescales of the HD signal during wakefulness and NREM sleep. As in ref. ^30^, we computed the square root of the rates to normalise for the variance in firing rates. Binned firing rates were smoothed with a Gaussian kernel of three bin s.d. (independent of absolute bin duration).

We then stacked together the binned spike trains from wakefulness and sampled 1% of all bins around DU in an interval of +/- 250ms, yielding a rate matrix ℝ ∈ ℝ^T×N^, where N is the number of neurons and T is the total number of time points. This rate matrix was then projected to a two-dimensional plane I ∈ ℝ^T×2^ using Isomap. The number of neighbours was set to 200. A subsampling of NREM bins was necessary due to the computationally intensive nature of this technique. This model of the fitted Isomap projection was then used to transform all the NREM binned rates. To ensure that the centre of the ring manifold lies at the origin, we subtracted the offset in the X- and Y-coordinates.

Ring radius was computed using the Euclidean distance of projected activity from the origin at each time point. The distribution of ring radius is typically bimodal, with large radii corresponding to wakefulness and UP states, and small radii corresponding to DOWN states. We empirically observed that the minima of this bimodality lies at one-third of the mean wake radius (**Supp Fig 1C**), which we used as a threshold for computing angular direction. Only epochs where radius exceeded the threshold at the previous UP state and post-DU were considered. Angular direction was computed using the element-wise arctangent.

We then selected all UP states larger than 500ms and computed angular directions for 0-150 ms post-DU and 150-300 ms post-DU, corresponding to the travelling UP state and control intervals, respectively. For both these intervals, we computed the cumulative circular variance (using SciPy.stats’ *circvar* function), starting with the first three bins and adding subsequent bins.

The diffusion coefficient is the average slope of the cumulative circular variance (using NumPy’s *gradient* function). Since the radius crossing the threshold during the travelling UP state in general varies between epochs, we computed the average slope of the cumulative circular variance plot in the interval of 35-150ms (35ms corresponding to the mode of the threshold crossing distribution).

### Delta peak detection

The wide-band signal was downsampled to 1.25 kHz and used as the LFP signal. The LFP was further downsampled to 250 Hz and filtered in the 0.5-4 Hz band using a second-order Butterworth filter. Peaks in the filtered signal during NREM epochs were detected using SciPy.signal’s *find_peaks* function. A peak was defined as a point that exceeded 3 s.d. of the filtered NREM delta band.

## Peri-Event Time Histograms (PETHs) and event-triggered average of LFP

Spikes from all putative excitatory neurons were binned in 5 ms bins and smoothed with a Gaussian kernel of 10 ms s.d. PETHs were computed on an interval of +/- 250 ms from DU. Isomap decomposition was performed on the PETHs of all sessions aligned to the DU transition (Fig 5C). The number of nearest neighbours was set to 50.

For the DU-triggered average of LFP, the wide-band signal was downsampled to 1.25 kHz and used as the LFP signal. The LFP was further downsampled to 625 Hz and filtered in the 70- 150 Hz band using a third-order Butterworth filter. This signal was smoothed with a Gaussian kernel of 32 ms s.d. and aligned to +/- 1 second around DU.

## Sequential Activation at UP onset

Following refs ^5,24^, to quantify when a given unit entered the UP state, we defined the UP onset delay as the time from the DU transition (up to 150 ms after DU) when the firing rate of the unit reaches 50% of its mean firing rate

Similarly, for quantifying sequential activation in the *γ*_h_ band, we defined the UP onset delay as the time from the DU transition where the derivative of *γ*_h_ reaches its maximal value (**Fig 2E**, inset). This quantity reflects the maximal change in *γ*_h_, corresponding to a DU transition.

We also computed the speed of the travelling UP state for both unit and LFP-based quantification. To do so, we used only those sessions that had significant correlations between UP onset delay and depth; while this means that sessions may not always be paired, it also ensures that our estimate of speed is not due to poor linear fits of the data.

## Wavelet Spectrogram

To compute the spectrogram around DU, the LFP was downsampled to 1.25 kHz. The spectrogram was computed on 100 logarithmically spaced frequencies going from 0.5 to 150 Hz. We constructed Morlet wavelets of 3 cycles centred around each frequency that were 4 s.d. in length on either side of the peak. These wavelets were then convolved with the downsampled LFP (using scipy.signal’s *fftconvolve* function). The absolute value of the result was taken to be the power of the frequency, which was z-scored +/- 1 second around the DU transition for each trial. We repeated this for all sessions and plotted the average spectrogram.

## Adapting Wilson/Cowan Model

The adapting Wilson-Cowan model was implemented as in Levenstein et al 2019, with the following modifications to extend the model to a 1D linear array.

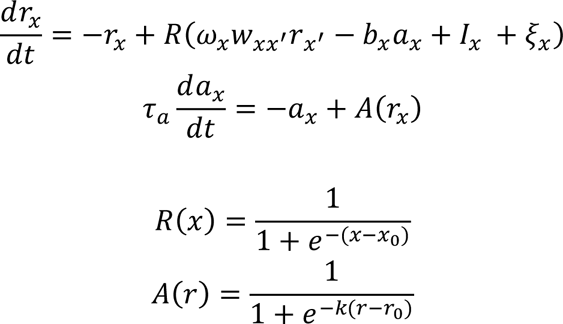

Where *r*_*x*_ and *a*_*x*_ are the time-varying rate and level of adaptation of the population at anatomical position *x*, *b*_*x*_, *I*_*x*_, and *ω*_*x*_ are parameters that represent are the strength of adaptation, input, and recurrence for the population at position *x*, *w*_*xx*_^′^ = *A* exp (−0.5((*x* − *x*/*s*)^2^) is the unit-normalized connection weight from the population at position *x*’ to *x*, and *ξ*_*x*_ = ζ(*t*) + ƞ_*x*_(*t*) is a combination of shared and independent sources of (Ornstein-Uhlenbeck) noise to the population at position *x* . Position was discretized to 100 units evenly spaced between *x* = 0 and *x* = 1.

To model gradients of different parameters (Figure 3B-D), we took either *b*, *I*, or *ω* to be a linear function of *x*, with the others constant. That is,

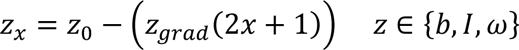

Where *z_grad_* is the magnitude of the parameter gradient and *z*_0_ is the mean value of the parameter over the entire population. Other parameters were held constant.

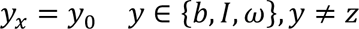

In the HDC data, the durations of DOWN states were shorter than UP states (mean_DOWN_ = 120.52 +/- 10 ms vs mean_UP_ = 596. 16 +/- 53.72 ms, computed across animals) and more regular (CV_DOWN_ = 0.63 +/- 0.04 vs CV_UP_ = 1.066 +/- 0.144, **Supp Fig 3A**), as previously reported for other cortical regions^23^. This indicates a dynamical regime in which a stable UP state is occasionally interrupted by noise-induced transitions to a transient DOWN state, during which the deactivation of adaptive processes returns the population to the UP state^23^. Parameter values were thus chosen as in Levenstein et al 2019 for an Excitable_UP_ regime and are presented in Table 1. For the parameter sweep (Supp Figure 3), the gradient magnitude for each parameter, *z_grad_*, and the connectivity width, *s*, were varied.

**Table 1:** Model parameters.

| Parameter | Value | Interpretation |
| --- | --- | --- |
| $\omega_0$ | 6.05 | Recurrent strength |
| $b_0$ | 1 | Adaptation strength |
| $I_0$ | -2.35 | Input strength |
| $\omega_{grad}$ | 0.075 | Recurrence gradient |
| $b_{grad}$ | 0.150 | Adaptation gradient |
| $I_{grad}$ | 0.085 | Input gradient |
| $\tau_a$ | 20 | Adaptation time constant |
| $x_0$ | 0 | Firing rate I/O curve midpoint |
| $k$ | 15 | Adaptation I/O curve steepness |
| $r_0$ | 0.5 | Adaptation I/O curve midpoint |
| $s$ | 0.1 | Recurrence width |

To model I_h_, the sign of adaptation’s effect on rate, *b*_0_, and its I/O curve, *k*, were reversed (Supp Figure 3E), and the input strength was decreased to compensate for the additive, rather than subtractive, effect of I_h_ (Table 2 for parameters). For clarity, the variable ℎ is used in place of *a*, though the form of the model is the same.

**Table 2:** Model parameters for I_h_.

| Parameter | Value | Interpretation |
| --- | --- | --- |
| $b_0$ | -0.75 | $I_h$ strength |
| $I_0$ | -3.2 | Input strength |
| $b_{grad}$ | 0.5 | $I_h$ gradient |
| $k$ | -30 | $I_h$ I/O curve steepness |
| $r_0$ | 0.3 | $I_h$ I/O curve midpoint |
| $s$ | 0.05 | Recurrence width |

The model was simulated for 20,000 timesteps using MATLAB’s *ode45* differential equation solver. DOWN states were then detected as times in which the mean rate over all positions fell below the dip in its bimodal distribution. To calculate the activation gradient, the mean rate was calculated for each unit aligned to DU and UD transitions (a PETH), and the time at which the PETH for each unit crossed *r* = 0.5 was found. The slope of the activation gradient was then taken to be the linear regression between the time of threshold crossing and the depth (*x*) of each unit. All model code is available at https://github.com/PeyracheLab/MehrotraLevenstein_2023.

## Slice preparation and whole-cell patch clamp recordings

Brain slices containing the HDC were prepared as previously described for other brain areas^107^. Briefly, mice were terminally anaesthetised with isoflurane and then decapitated. The brain was rapidly dissected into ice-cold, semi-frozen sucrose-ACSF (in mM: 87 NaCl, 2.5 KCl, 25 NaHCO_3_, 1.25 NaH_2_PO_4_, 25 glucose, 75 sucrose, 7 MgCl_2_, 0.5 CaCl_2_, 1 Na-Ascorbate, 1 Na-Pyruvate) bubbled with carbogen (95% O_2_/5%CO_2_). 300 μm thick coronal brain slices were cut on a vibratome (VT1200s, Leica, Germany) from the posterior portion of the cerebral cortex containing the dorsal HDC. Slices were then transferred to a submerged storage chamber containing sucrose-ACSF at 35°C for 30 min and subsequently at room temperature.

For recording, slices were transferred to a submerged recording chamber which was perfused with carbogenated recording ACSF (in mM: 125 NaCl, 2.5 KCl, 25 NaHCO_3_, 1.25 NaH_2_PO_4_, 25 glucose, 1 MgCl_2_, 2 CaCl_2_) at a rate of 6-8 ml/min at 31 ± 1°C by an inline heater (Scientifica, UK). Slices were visualized under Köhler illumination by means of an upright microscope (Slicescope, Scientifica, UK), equipped with a 4x air-immersion (N.A. 0.13; Olympus) and 40x water-immersion (N.A. 0.8; Olympus) objective lenses. Whole-cell patch-clamp recordings were performed using a MultiClamp 700B amplifier (Molecular Devices, CA, USA). Intracellular recording electrodes were pulled from borosilicate glass capillaries (1.5 mm outer/0.86 mm inner diameter, Harvard Apparatus, UK) on a horizontal electrode puller (P-1000, Sutter Instruments, CA, USA). When filled with intracellular solution (in mM: in mM: 142 K-gluconate, 4 KCl, 0.5 EGTA, 10 HEPES, 2 MgCl_2_, 2 Na_2_-ATP, 0.3 Na_2_-GTP, 1 Na_2_-Phosphocreatine, 2.7 Biocytin; pH = 7.4, osmolarity: 290-305 mOsm) a pipette resistance of 4-6 MΩ was achieved. Unless otherwise stated, all recordings were performed in current clamp from resting membrane potential (V_M_). For all recordings, the bridge was balanced following pipette-capacitance compensation in current-clamp. Signals were filtered online at 10 kHz using the built in 2-pole Bessel filter of the amplifier and digitized at 20 kHz (Digidata 1550B, Axon Instruments, USA) using pClamp 10 (Molecular Devices, CA, USA). Data was analysed offline using the open source Stimfit software package^108^ (http://www.stimfit.org). The liquid junction potential was measured as 12 mV in the recording configuration, but not adjusted.

Cells were selected for recording in layer 3 of the HDC directly below layer 2 and had round to ovoid somata with prominent apical dendrites extending towards layer 1. We identified the HDC/retrosplenial cortex border under IR-DIC illumination, then recorded cells from this point to 1200 µm ventral. Recordings were interleaved such that both dorsal, ventral, and medial cells were recorded in turn, with no one cell type always recorded first or last. After obtaining a whole-cell recording, the intrinsic physiology of recorded neurons was characterised in current-clamp mode from resting membrane potential. Upon break-through, 30x -10 pA steps were immediately applied to obtain resting membrane potential, input resistance, and membrane time-constant. Following this, we applied 3 runs of 500 ms hyper-to depolarising current steps (-500 to +500 pA, 100 pA steps). Following characterisation, each cell was re-sealed with outside-out patches, then further cells were recorded in each slice. For the last cell in each slice, 20 µM ZD-7,288 (Tocris, UK) was applied to the bath for 5 minutes, then intrinsic characterisation performed again. I_h_-mediated voltage sag was estimated from -500 pA current steps, as the difference between the initial peak hyperpolarization and the steady state voltage response. This was normalised to the peak response to account for cell-to-cell resistance differences. Action potential properties were measured from the first spike elicited at rheobase current. All measurements are the average of at least 3 repetitions/cell.

## Post-hoc visualisation and imaging

Following recording and sealing of the final cell of each slice, tissue was fixed overnight in 4% paraformaldehyde (in 0.1 M Phosphate Buffer (PB), pH 7.4) at 4°C. To visualise neurons, the slices were washed in 0.1 M Phosphate Buffered Saline (PBS), and then incubated overnight at 4°C with 2 µg/ml of fluorescent-conjugated streptavidin (AlexaFluor633, Invitrogen, UK), which was diluted in 0.1M Phosphate Buffered Saline (PBS) and 0.5% Triton-X 100. Slices were then washed liberally in PBS, then PB, and mounted on glass slides with fluorescent protectant mounting medium (FluoMount-G, Southern Biotech, USA). Slices were imaged on an inverted confocal microscope (SP8, Olympus, Japan) to generate low-power (20x optical magnification) overviews of recorded neurons. Based on these overviews, the position of recorded neurons was calculated, relative to the dorsal pole of the HDC.

## Statistics

All analyses were conducted using software custom-written in Python 3.10, using the Pynapple package^109^, except for the *ex vivo* data that was analysed using GraphPad Prism. Unless otherwise specified, statistical comparisons were performed with Kendall τ rank correlation, Mann-Whitney U test, Wilcoxon Signed-Rank Test, or Pearson’s tests, where applicable.

## Acknowledgements

We would like to thank Lynda Mainville for technical support. We are thankful to the members of Peyrache laboratory for comments on an earlier version of the manuscript.

## Funding

Canadian Research Chair in Systems Neuroscience (AP), CIHR Project Grant 155957 and 180330 (AP), NSERC Discovery Grant RGPIN-2018-04600 (AP), Canada-Israel Health Research Initiative, jointly funded by the Canadian Institutes of Health Research, the Israel Science Foundation, the International Development Research Centre, Canada and the Azrieli Foundation 108877-001 (AP), Sir Henry Wellcome Fellowship 206491/Z/17/Z (AJD), Embo Long-Term Postdoctoral Fellowship ALTF 382-2017 (AJD), Doctoral training award from the Fonds de Recherche du Québec, Santé (DM), UNIQUE Postdoctoral Excellence Scholarship (DL) and the Richard and Edith Strauss Postdoctoral Fellowship in Medicine (DL).

## Author contributions

Conceptualization: DM, DL, AJD, AP; Methodology: DM, DL, AJD, AP; Software: DM, DL, AJD, AP; Validation: DM, DL, AP; Formal analysis: DM, DL, AJD, SAB, AK, AP; Investigation: DM, DL, AJD, SSC, SAB, AK; Resources: AP; Data Curation: DM, DL, AJD, SAB, AK; Writing - Original Draft: DM, DL, AP; Writing - Review & Editing: DM, DL, AJD, AP; Visualization: DM, DL, AJD, AP; Supervision: AP; Project administration: AP; Funding acquisition: AP, AJD

## Competing interests

Authors declare that they have no competing interests.

## Supplementary Information

**Supplementary Figure 1, related to figure 1:**
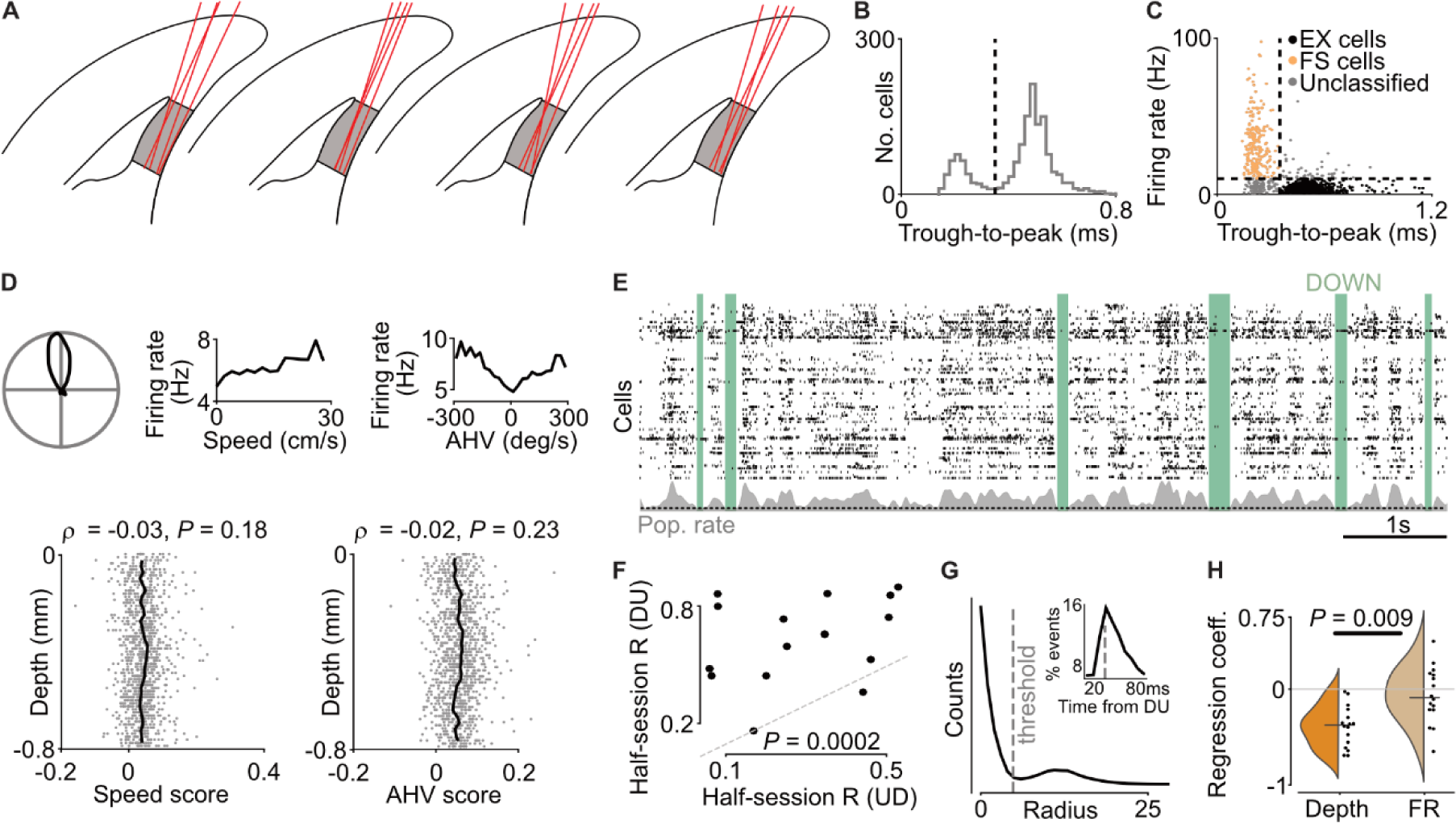
**A)** schematic of estimated probe trajectories as they were lowered towards HDC. Probes were positioned so that the top channels corresponded to the dorsal edge of HDC. **B)** Histogram showing bimodal distribution of trough-to-peak duration of all recorded cells. Dashed line, threshold for classifying EX cells. **C)** Scatterplot showing classification of EX cells based on through-to-peak and mean firing rate. **D)** Top, example of an EX cell tuned to HD, linear speed, and angular head velocity (AHV). Bottom, quantification of linear speed and AHV tuning in EX cells as a function of HDC depth. **E)** An 8 s sample of HDC spiking is shown (black), along with the population rate (grey) and the detected DOWN states (green). **F)** Half-session correlation for UP state onset and the UD transition suggests that the UP state onset is overall more stable than the UD transition (Wilcoxon signed-rank test, W = 4, n = 16, *p* = 0.0002). Grey dashed line indicates the unity line. **G)** Distribution of Isomap ring radii for an example session. The distribution is bimodal, consisting of small radii (observed during NREM but not wakefulness) and larger radii (observed during wakefulness and UP states). The threshold was computed as one-third the mean wake radius, which empirically corresponds well with the local minima of the distribution. Inset: The distribution of UP onset delays where the radius crosses the threshold described previously. The grey line indicates the cutoff used for the diffusion constant analysis, i.e., 35 ms. **H)** The regression coefficient of UP onset delay as a function of anatomical position (“depth”) and NREM firing rate (“FR”), each observation represents one animal. The coefficients for anatomical position explain the UP onset delay better than the firing rate (Wilcoxon signed-rank test, W = 19, n = 16, *p* = 0.009), and are significantly different from 0 (One-sample Wilcoxon test, W = 0, n = 16, *p* = 3.052 x 10^-5^).

**Supplementary Figure 2, related to figure 2:**
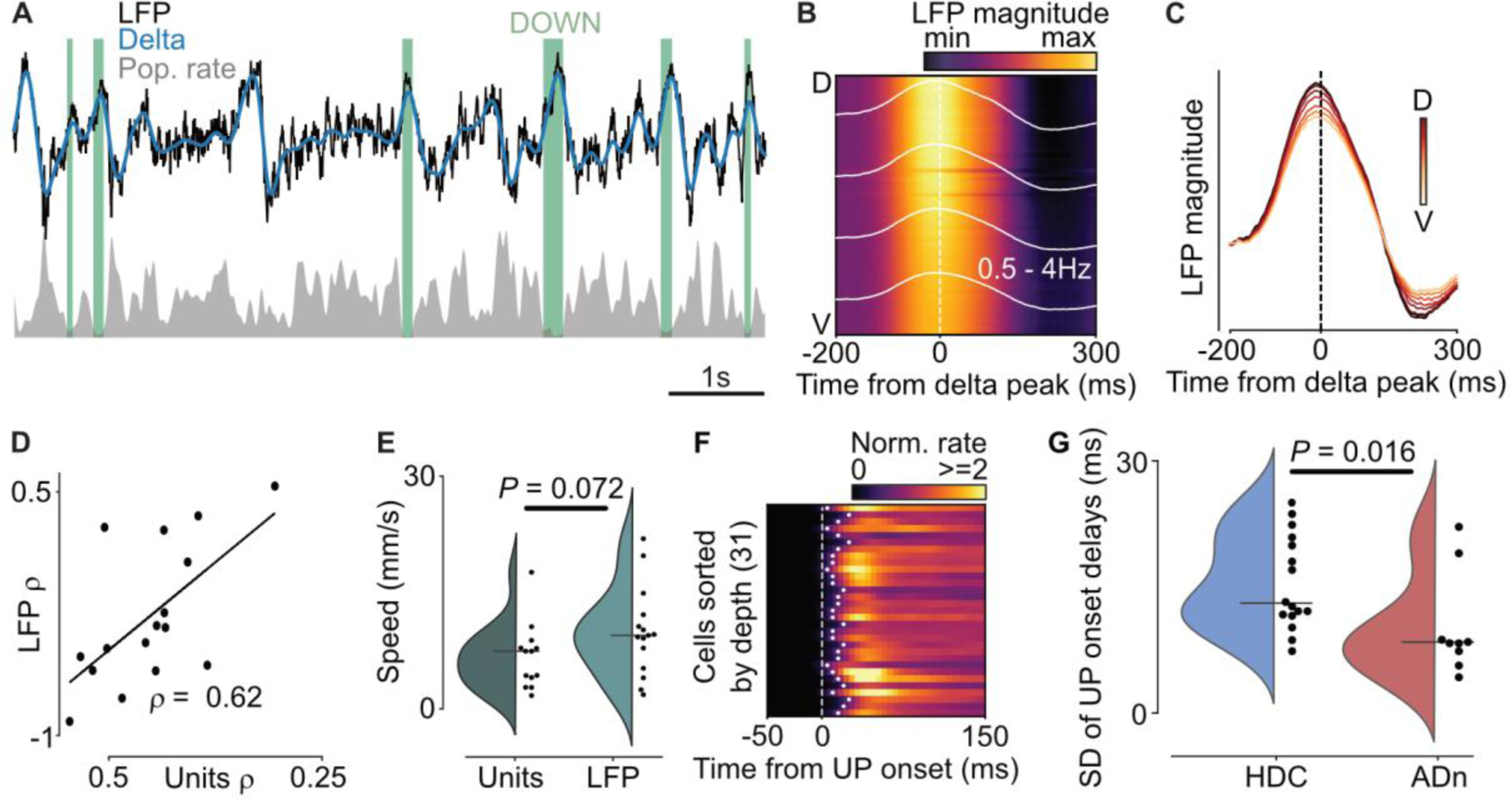
**A)** An 8s sample of HDC LFP is shown (black), along with the delta band (0.5 - 4 Hz) activity, and population rate (grey). Detected DOWN states based on population activity are shown in green. Note that the delta peak is related to a drop in the population activity. **B)** Example epoch showing broadband LFP magnitude aligned to the delta peak (dashed line). The white line indicates the delta-band LFP for 4 example channels. Note that the LFP is synchronous along the dorsoventral axis. **C)** Delta peak aligned broadband LFP for 8 example channels (coloured by anatomical position). **D)** Correlation between UP onset delay and anatomical depth, as computed by units (X-axis), versus LFP (Y-axis) show an agreement between the measures (Pearson R = 0.62, n = 16, *p* = 0.01). **E)** Speed of the travelling UP onset, measured as the slope of the UP onset delay v/s depth computed using units and LFP yield similar values of the speed of travel (Mann-Whitney U test, U = 58, n = (13, 15), *p* = 0.072). The speed was computed only for sessions with significant correlations between UP onset delay and anatomical depth. **F)** Representative PETH of ADN cells aligned to the DU transition. White dots indicate the bin where the firing rate exceeded 50% of the mean firing rate. Note the synchronous reinstatement of the UP state in the dorsoventral direction. **G)** Distribution of the range of UP onset delays computed over all sessions of recorded ADN and HDC neurons indicate that the sequential activation of UP onset delays is likely a local phenomenon arising in the HDC and is not driven by synaptic inputs from ADN (Mann-Whitney U test, U = 29, n = (16, 9), *p* = 0.016).

**Supplementary Figure 3, related to figure 3:**
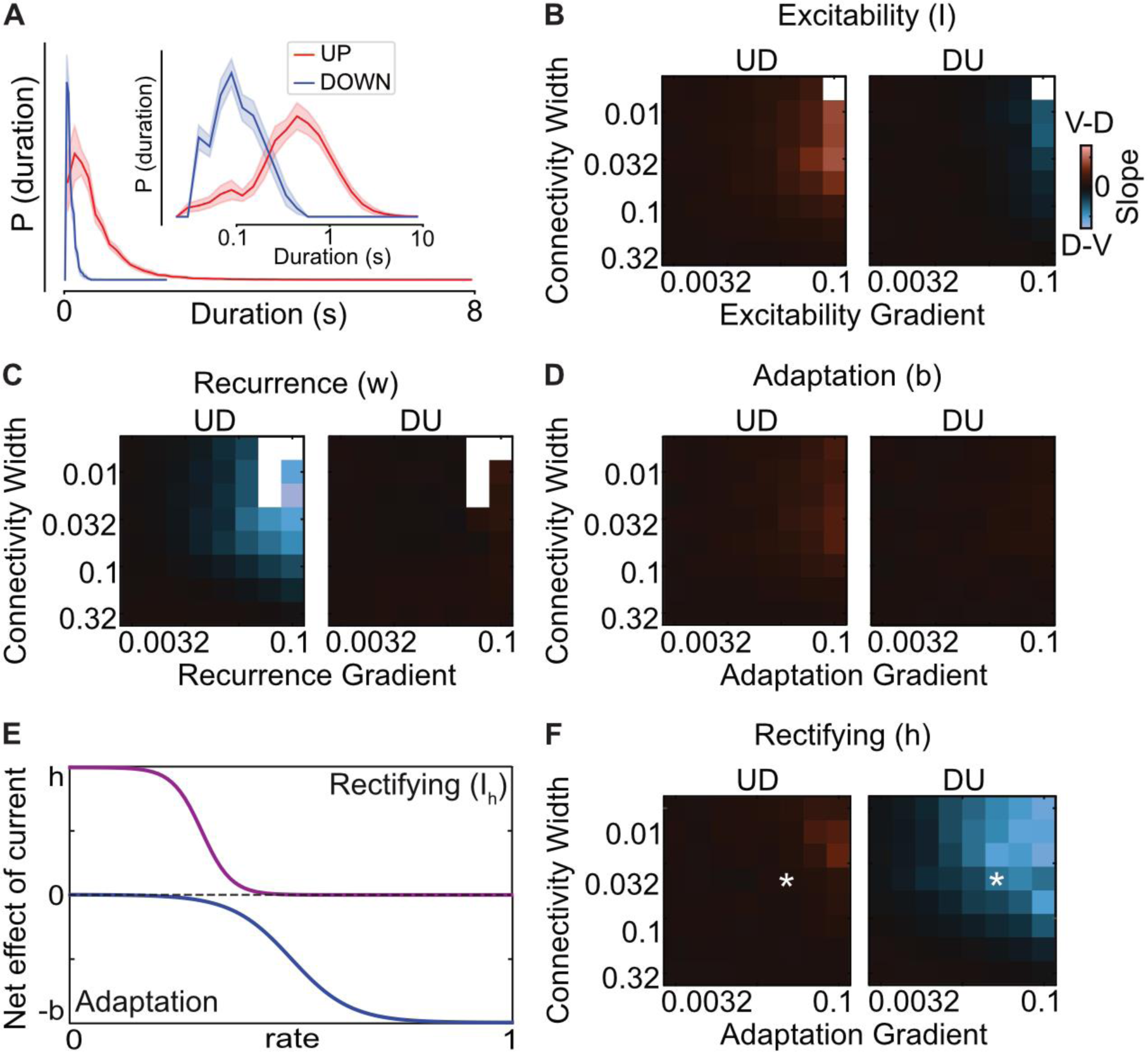
**A)** Distribution of UP/DOWN state duration in an example recording, on a linear and logarithmic scale (inset). **B-D)** Parameter sweep of the adapting Wilson-Cowan model with a gradient of excitability **(B)**, recurrence **(C)**, and adaptation **(D)** are unable to replicate experimental observations. **E)** Motivation behind the rectifying current as an inverse-adapting current; traditional adapting currents have no effect at low rates and a subtractive effect at high rates, while the rectifying current has no effect at high rates and an additive effect at high rates. This is equivalent to merely shifting the activation profile of the adapting current along the positive Y-axis. **F)** Parameter sweep of the model with rectifying currents shows that it can replicate experimental findings robustly over a range of parameter values.

**Supplementary Figure 4, related to figure 4:**
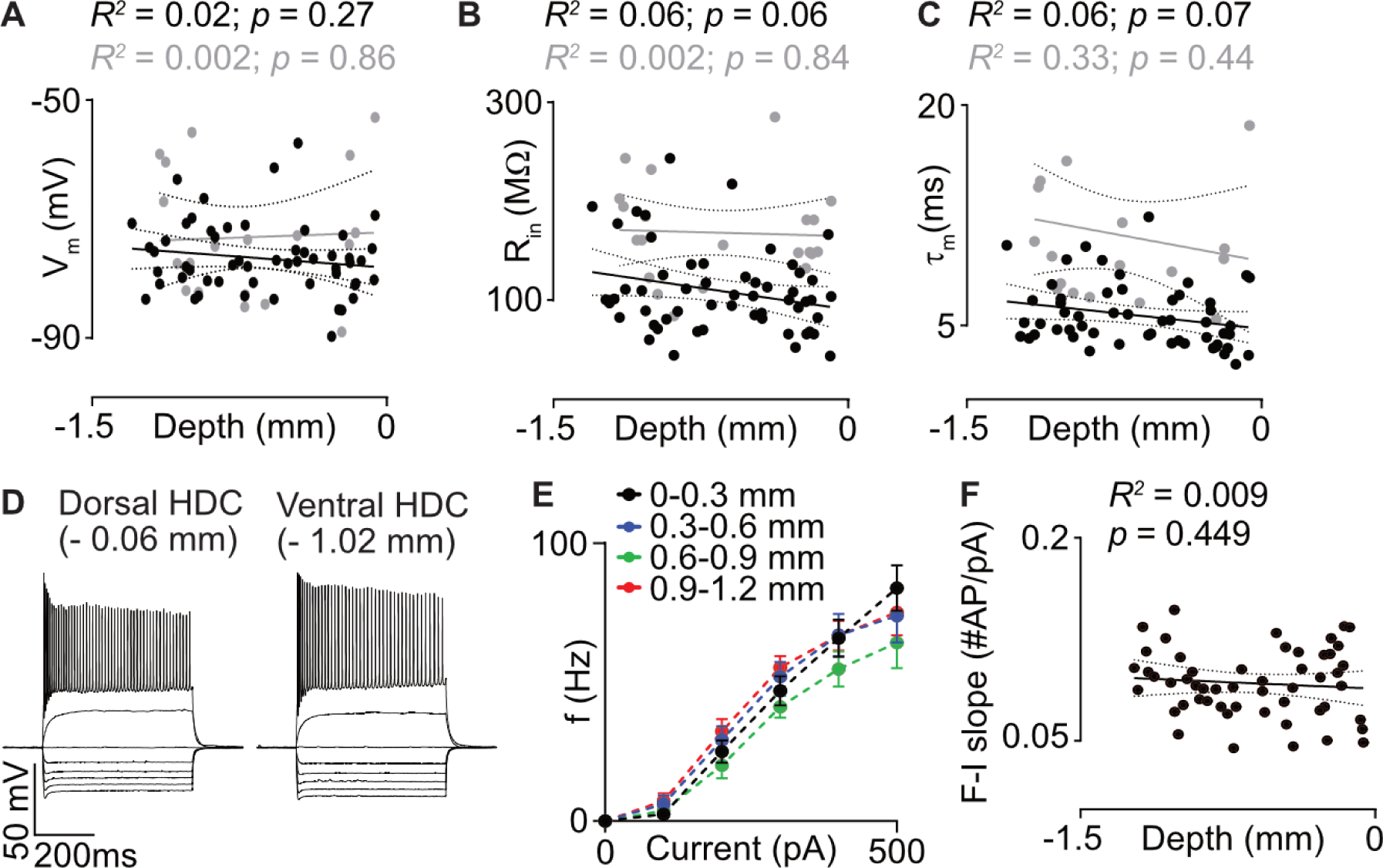
**A-C)** Passive properties in the HDC show no relationship with anatomical position in control conditions (black, n = 59 cells from 11 mice), or when treated with 20μM ZD-7288 (grey, n = 19 cells from 11 mice), as observed for **A)** resting membrane potential (V_m_), **B)** input resistance (R_in_) and **C)** membrane time constant (τ_m_). **D)** Representative voltage response to current clamp for cells in dorsal and ventral HDC (Currents used here are -500, -400, -300, -200, -100, 0, +100, +500 pA, all 500 ms duration). Note the increased voltage sag in the dorsal HDC traces. **E)** F-I curves for cells at different dorsoventral positions in the HDC show similar characteristics. **F)** Slope of the F-I curve does not vary with anatomical position in the HDC.

**Supplementary figure 5, related to figure 5:**
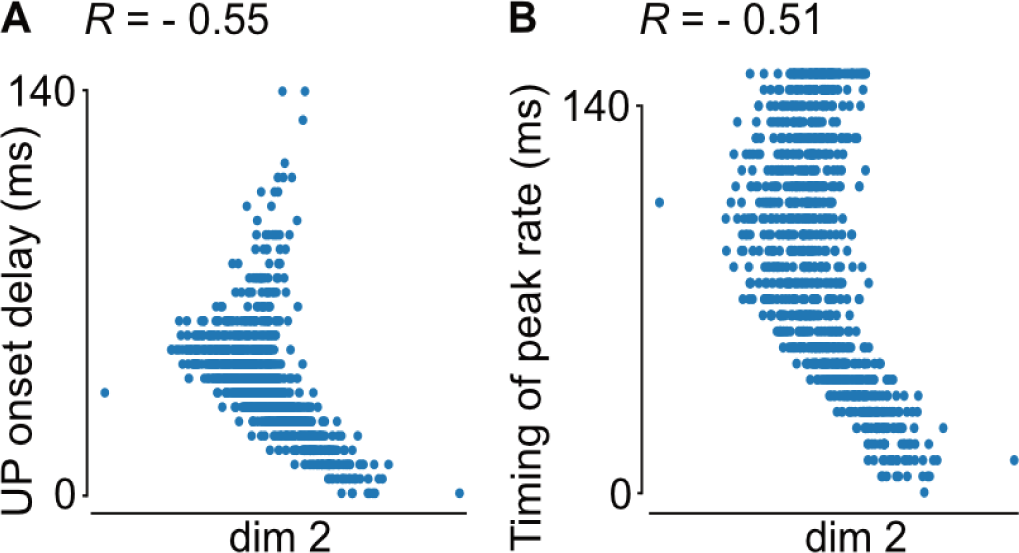
The second dimension of the DU-PETH Isomap projection is correlated with **A)** UP onset delay (Pearson R = -0.55, n = 1096, *p* = 3.39 x 10^-89^) and **B)** timing of peak firing rate (Pearson R = -0.51, n = 1096, *p* = 1.64 x 10^-73^), indicating that the second dimension is better correlated with UP onset delay than with PMR. Taken together, PMR and UP onset delay form the axes of maximal variance in the PETH Isomap.

## Notes

### Competing Interest Statement

The authors have declared no competing interest.

